# Estimating body volumes and surface areas of animals from cross-sections

**DOI:** 10.1101/2023.10.13.562315

**Authors:** Ruizhe Jackevan Zhao

## Abstract

Body mass and surface area are among the most important biological properties, but such information are lacking for some extant organisms and all extinct species. Numerous methods have been developed for body size estimation for this reason. There are two main categories of mass-estimating methods: volumetric-density approaches and extant-scaling approaches. In this paper, a new 2D volumetric-density approach named cross-sectional method is presented. Cross-sectional method integrates biological cross-sections to obtain volume and surface area accurately. Unlike all previous 2D methods, cross-sectional method processes true cross-sectional profiles directly rather than approximating. Cross-sectional method also has the advantage over others that it can deal with objects with gradually changing cross-sections. It generates very accurate results, with errors always lower than 2% in all cases tested.

## Introduction

Body mass and surface area are related to many biological properties such as physiology, ecology and evolution (e.g, [1, 2, 3, 4]). However, body masses are unavailable for many large extant animals and all extinct species. Surface area information is also lacking because it can not be measured directly. To solve this problem, previous researchers have developed numerous methods for body size estimation.

In general, there are two categories of approaches for body mass estimation: extant-scaling methods and volumetric-density methods [5]. Extant-scaling methods utilize skeletal measurements as proxies and discover their relationships with body mass using regression [6, 7]. A classic and universally applied example of extant-scaling approaches is the equation for quadruped mass based on humeral and femoral circumference [8].

The priciple of volumetric-density methods is to obtain the body volume first, then an overall density is assigned to transform volume into mass [9, 10, 11]. They have a much longer history than extant-scaling methods, and numerous approaches have been developed over the past century. The earliest volumetirc-density methods are based on physical models. Gregory [12] soaked a *Brontosaurus* model in water and acquired its volume, then he scaled the result to get the true size.

Some mathematical methods were developed later to calculate volume and surface area from two-dimensional profiles. The first 2D volumetric-density method, Graphic Double Integration (GDI), was invented and introduced by Jerison [13]. Henderson [10] developed a more rigorous math method called mathematical slicing to calculate volume and center of mass. Both GDI and mathematical slicing partition animals into several parts (known as ‘slabs’ in [10]) and treat them as frustums with elliptical bases. Seebacher [14] invented a polynomial method, which uses polynomials and curve equations to simulate body outlines and cross-sections respectively. Motani [11] argued that superellipses are better approximations of biological cross-sections rather than ellipses and developed the first version of Paleomass. The latest study on 2D volumetric-density methods is the new version of Paleomass implemented in R [15].

With the rise of computer technology, 3D modeling has been widely applied in animal reconstructions (e.g., [16, 17, 18]). The first step of 3D reconstruction of extinct vertebrates is to obtain the skeleton, which can be converted from photographs or 3D scans, then soft tissue is added to the skeleton [16, 5]. During this process, errors and subjectivity can not be avoided [5]. Sellers et al [19] invented convex hull method, which generates minimum convex hulls to envelope the skeleton and adjusts the amount of soft tissue based on extant mammals. Convex hull method can reduce the errors introduced during soft tissue reconstructions, but it has the disadvantage that a large quantity of extant organisms are required as samples [15]. Comparing with 2D approaches, 3D modeling requires proficient use of 3D software and is more time-consuming, so there is still demand for developing 2D methods.

In this paper, a new 2D volumetric-density approach named ‘cross-sectional method’ is presented. Instead of assuming elliptical or superelliptical approximations, it processes gradient cross-sectional profiles directly and is able to handle any shapes. It produces body volume and surface area calculation results with high accuracies.

## Materials and Methods

### Data Collection

The side view (or dorsal/ventral view) outline of the studied animal is collected first, by drawing along the profile from photos, precise life reconstructions or orthogonal projections of 3D models. Biological structures like flukes, limbs and horns are then separated from the main body (Fig. 1A). Their volumes and surface areas will be calculated independently using the same method as in the main body part (see below).

**Figure 1:**
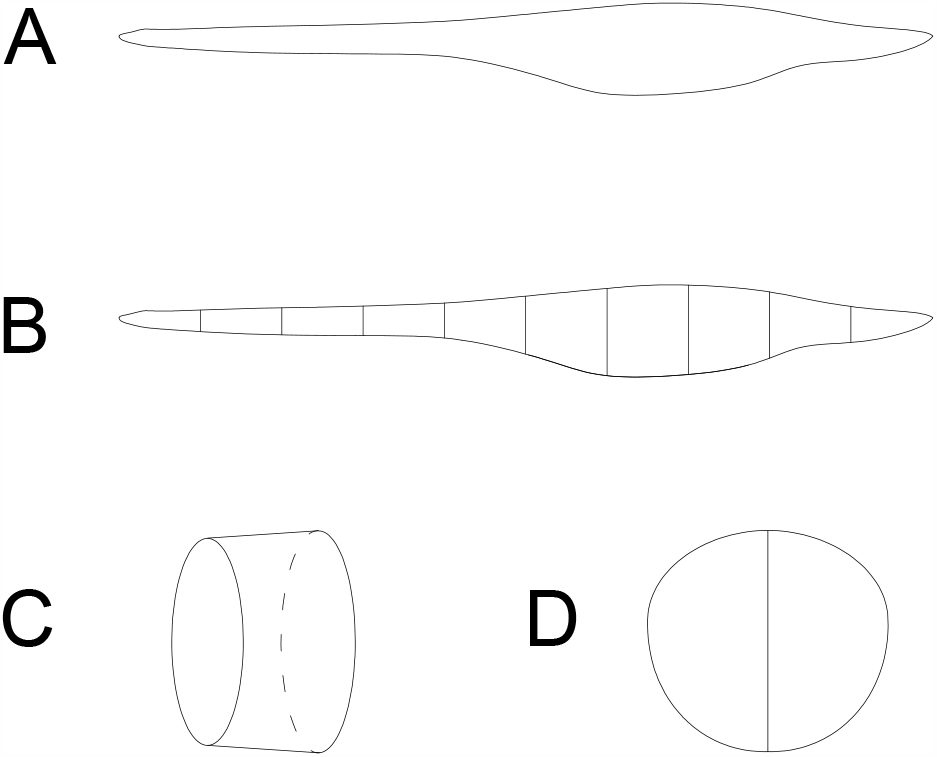
Illustrative process of data collection. (A) Collect the side view (or dorsal/ventral view) outline; (B) Slice the profile into slabs; (C) Illustration of a slab; (D) An example of cross-section with an identity segment (vertical).

The terms ‘slab’ and ‘subslab’ used by Hederson [10] are inherited here. After the outline is obtained, the animal’s profile is equally partitioned into several slabs using parallel lines (Fig. 1B, C). The accuracy of the calculation increases together with the number of slabs. The portions of parallel lines truncated by the profile (i.e., maximum heights in side views or maximum widths in dorsal/ventral views) are defined here as ‘identity segments’ (Fig. 1D).

After partitioning, each slab (except the first and last one) can be viewed as a frustum with parallel bases, which are probably different in shape (Fig. 2). The slabs at two ends of the animal’s sagittal axis can be viewed as cones with irregular-shaped bases.

**Figure 2:**
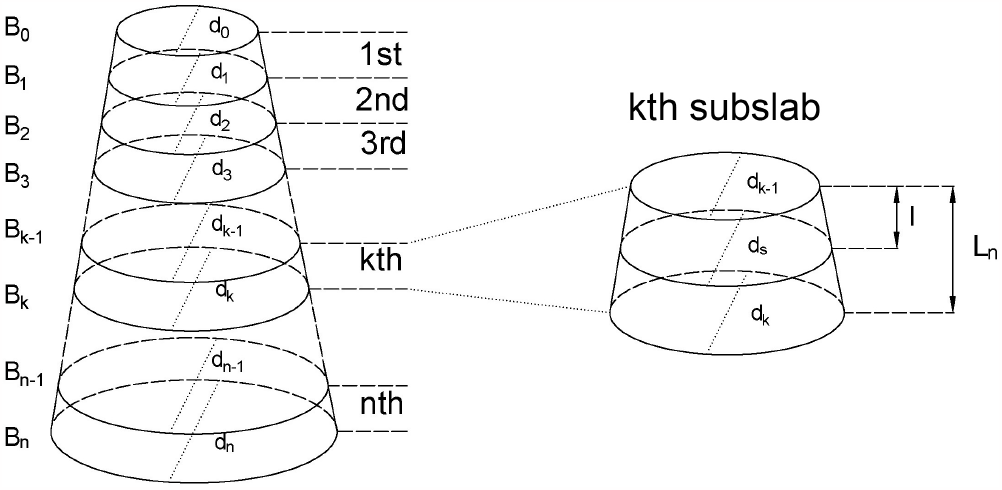
Illustration of Slab and Subslab.

The next step is to collect the profiles of bases in each slab, which are originally body cross-sections of the studied animal (Fig. 1D). Then the area and circumference of each cross-section are acquired using image processing software.

### Body Volume Calculation

Consider a slab of which the two parallel bases are different in shape (Fig. 2). The two bases both have an identity segment (denoted by *d*_0_ and *d*_*n*_ respectively) as proxies for their areas (denoted by *S*_0_ and *S*_*n*_ respectively). The ratio of *S* to *d*^2^ is defined and denoted by *φ*, i.e.,

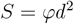

Then slice the slab equally into *n* subslabs with all the bases parallel to each other, where *n* is a large positive integer. The upper base and lower base of the *k*th subslab are denoted by *B*_*k−*1_ and *B*_*k*_. The parameters (as defined above) of the lower base of the *k*th subslab are denoted by *d*_*k*_, *S*_*k*_, and *φ*_*k*_ respectively. The height of the slab is denoted by *L*, and the height of each subslab is *L*_*n*_. Assume that *φ*_*k*_ follows a linear relationship from *φ*_0_ to *φ*_*n*_, then

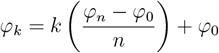

Now consider the volume of the *k*th subslab. Lengths of identity segments (*d*) can not be simply assumed to increase or decrease linearly, because maximum body heights/widths along an animal’s sagittal axis often show irregular undulation. However, in calculus linearity is often used to approximate non-linearity at very small scales. Technically, it’s impossible to slice a real object into infinite partitions. Since *n* is a large number, it can also be assumed that within each subslab *d* also follows a linear relationship. Then for any cross-section (denoted by *B*_*s*_) in the *k*th subslab parallel to the bases *B*_*k−*1_ and *B*_*k*_, it holds that

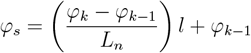

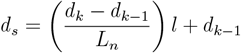

where *l* is the distance from *B*_*s*_ to *B*_*k−*1_.Then let

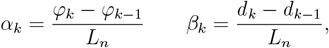

The area of cross-section *B*_*s*_ can be calculated by

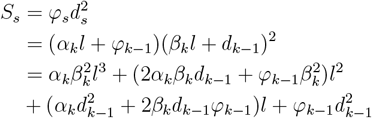

Then the volume of *k*th subslab is

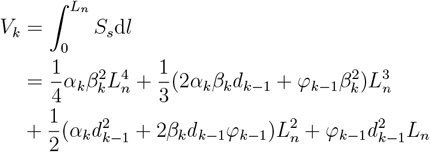

In particular, if *φ* is a constant (denoted by Φ), then *α*_*k*_ = 0 and

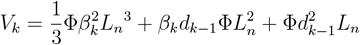

The total volume of the slab is

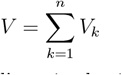

The two slabs at both ends of the animal’s sagittal axis are processed as slabs with constant Φ, others are regarded to possess gradually changing cross-sections. The total main body volume can be acquired by summing the volumes of all the slabs. The volumes of structures separated (e.g., fins, limbs) from the main body are calculated using the same method.

Previous studies assigned different overall body densities to different animals (e.g., [20, 21]), but a discussion of density variation among broad taxonomic clades is beyond the scope of this study. In this paper, all animal models are treated as solid objects, with densities not assigned (i.e., only the overall volumes are studied). Future scholars can easily acquire body mass estimations by assigning cavity sizes and densities to volumes calculated using the cross-sectional method.

### Body Surface Area Calculation

Similar method is applied to calculate the surface area (Fig. 2). All parameters defined in volume calculation except *φ* are inherited here. The circumferences of the upper base and lower base of the slab are denoted by *C*_0_ and *C*_*n*_. The ratio of *C* to *d* is denoted by *ψ*, i.e.,

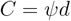

The parameters (as defined above) of the lower base of the *k*th subslab are denoted by *d*_*k*_, *C*_*k*_, and *ψ*_*k*_. Assume that *ψ*_*k*_ follows a linear relationship from *ψ*_0_ to *ψ*_*n*_, then it holds that

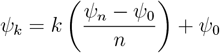

After slicing the slab equally into *n* subslabs, linearity is used to approximate non-linearity at a very small scale:

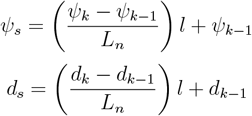

where *l* is the distance from *B*_*s*_ to *B*_*k−*1_. Then let

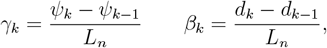

The circumference of cross-section *B*_*s*_ can be calculated by

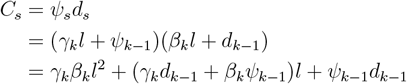

Then the lateral surface area of *k*th subslab is

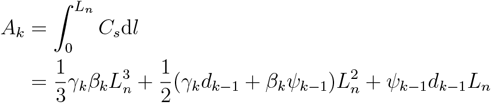

In particular, if *ψ* is a constant (denoted by Ψ), then *γ*_*k*_ = 0 and

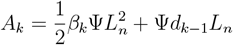

The total lateral surface area of the slab is

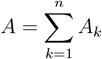

The two slabs at both ends of the animal’s sagittal axis are processed as slabs with constant Ψ, others are regarded to possess gradually changing cross-sections. The surface area of the main body is calculated by summing the lateral areas of all slabs. The surface areas of structures separated (e.g., fins, limbs) from the main body are calculated using the same method.

### Validation and Comparison

To test the accuracy of cross-sectional method, two tests are carried out. In both tests, the volumes and surface areas of 3D models are first obtained, then the calculated results based on 2D methods are compared with the true values to validate their accuracy. Only models precisely reproduced from museum mounts, life photos or 3D scans are used for validation (see [22, 23]; and http://digitallife3d.org/). To further evaluate the performance of cross-sectional method, GDI and Paleomass are included as representative methods for comparison, which approximate biological cross-sections with ellipses or superellipses [9, 15]. To ensure that the three methods can be compared in a same framework, thirteen 3D models of extinct or extant aquatic species are used.

Before the tests, protruding structures like limbs, flukes and fins are separated from the main body. Different structures from a same model may be used in different tests (see below).

The first test is based on 10 secondarily aquatic tetrapod models (5 plesiosaurs, 3 ichthyosaurs and 2 cetaceans). To test the performances of the three methods when handling main bodies, flippers and fins of these models are removed. The main body of each model has rounded or oval cross-sections, which can be well-approximated by ellipses or superellipses.

In the second test, biological structures with irregular cross-sections (see discussion) are used. Such objects include fins and limbs of aquatic animals. The main body of an Atlantic sturgeon (*Acipenser oxyrhynchus oxyrhynchus*) model and a hawksbill turtle (*Eretmochelys imbricata*) model are also included.

In GDI, each object is first equally sliced into 10 slabs, then the volume is calculated using the formula proposed in [9] after necessary measurements are made. Paleomass is performed using the corresponding package in R [15]. The four fin/flipper samples in the second test are treated as foils and others are treated as main bodies (see [15] for detailed methods). k-value range 2-2.3 and 1.6-2.4 are used for the two tests respectively. The former is the range suitable for modern cetaceans, and the latter successfully bracketed all aquatic species tested in [15]. In cross-sectional method, each object is equally sliced into 10 slabs and each slab is further sliced into 10 subslabs, then the volume and surface area are calculated after parameters of the bases in each subslab are obtained.

After the calculation, the amounts of error generated by different methods are compared. Error is defined as

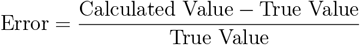

When the calculation underestimates the true value, the error is negative; when overestimating, the error is postive. Afterwards the mean error is calculated as:

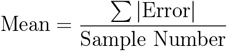

### Software Application

In this study, all the 3D models are first processed in Rhino 7. Each model is separated using *WireCut* command, then the true volume and surface area of the selected part are acquired using *Volume* and *Area* command respectively. Side view and dorsal/ventral view images of the separated models are obtained with *Make2D* command. To generate the cross-sections needed in cross-sectional method, *ClippingPlane* command is used.

Two dimensional images are then imported into AutoCAD 2020, where they are sliced into slabs or subslabs using *Arrayrect* and *Trim* commands. Measurements of each slab or subslab are taken and exported into Excel with *Dataextraction* command. In crosssectional method, areas, circumferences and lengths of identity segments of the bases in each subslab are first measured with *Measuregeom* command, then the parameters (*φ, ψ*) are calculated with the calculator implemented in AutoCAD. The calculation of GDI and cross-sectional method is finally performed in Excel.

Both Rhino and AutoCAD are industrial software with a high precision. They have already been applied in previous studies for body size estimation of animals and proved to have good performances (e.g., [10, 24]).

Paleomass implemented in R requires bitmaps [15], so the two-dimensional images are exported from AutoCAD as PNGs. Each PNG is set to possess 1680 *×* 1280 pixels since Motani [11] suggested that Paleomass has better performance when handling images with high resolution. They are then imported into PhotoShop 2020 for dyeing. Afterwards the processed images are imported into R 4.1.3, where the final calculation takes place.

## Results

The results of the first test and second test are listed in Table 1 and Table 2 respectively. Main bodies that aren’t bracketed by selected k-value range successfully in Paleomass are marked with red color.

**Table 1:** Errors in the first test.

|  | Volume |  |  | Surface Area |  |  |
| --- | --- | --- | --- | --- | --- | --- |
| Model | GDI | Paleomass | CS | GDI | Paleomass | CS |
| *Dolichorhynchops* | −3.51% | 18.05% | −0.02% | −4.07% | 11.65% | −1.88% |
| *Hydrotherosaurus* | −5.33% | −0.48% | −0.54% | −4.30% | −0.24% | −1.17% |
| *Liopleurodon* | −0.16% | 4.86% | −0.02% | −2.81% | 2.20% | −1.94% |
| *Peloneustes* | −3.11% | −1.38% | 1.00% | −6.19% | −0.99% | −2.44% |
| *Thalassomedon* | −5.53% | 6.64% | −0.19% | −4.34% | 5.12% | −0.53% |
| *Ophthalmosaurus* | −3.90% | −1.29% | 0.16% | −3.28% | −0.28% | −2.04% |
| *Stenopterygius* | −2.28% | −0.07% | −1.82% | −2.63% | 0.41% | −2.35% |
| *Temnodontosaurus* | −2.46% | 4.33% | 0.12% | −2.70% | 2.04% | −0.75% |
| *Megaptera* | 1.81% | 6.41% | 0.25% | −1.65% | 2.54% | −1.52% |
| *Orcinus* | −2.88% | 3.13% | −0.16% | −0.44% | 4.34% | 0.79% |
| Mean | 3.10% | 4.66% | 0.43% | 3.24% | 2.98% | 1.54% |

**Table 2:** Errors in the second test.

|  | Volume |  |  | Surface Area |  |  |
| --- | --- | --- | --- | --- | --- | --- |
| Model | GDI | Paleomass | CS | GDI | Paleomass | CS |
| *Liopleurodon* flipper | 14.29% | 30.21% | −0.25% | 6.22% | 6.56% | −0.06% |
| *Orcinus* dorsal fin | 12.92% | −33.14% | −0.27% | 7.66% | −8.21% | −1.10% |
| *Ophthalmosaurus* tail fin | 12.26% | 15.44% | −0.41% | 6.18% | 0.74% | −1.40% |
| *Mobula* pectoral fin | 16.68% | −85.02% | −0.48% | 6.26% | −11.31% | −1.93% |
| *Acipenser* main body | 12.51% | 39.92% | −0.28% | −0.88% | 25.09% | −1.49% |
| *Eretmochelys* main body | 17.45% | 6.19% | 0.10% | 4.85% | −0.13% | −1.38% |
| Mean | 14.35% | 34.99% | 0.30% | 5.34% | 8.67% | 1.23% |

In the first test, all the three methods validated show good performances, with errors lower than 5% than by average. This corroborates the validity of 2D volumetric-density methods when handling animals with rounded or oval cross-sections, as demonstrated in previous studies [10, 15]. In both volume and surface area calculation, cross-sectional method shows slightly higher accuracies than others, proving its effectiveness in estimating volume and surface area.

In the second test, the errors of GDI and Paleomass increase significantly. This result indicates that an elliptical approximation, as assumed in GDI, are not suitable for all biological cross-sections. Paleomass treats the four fin/flipper samples as foils, which are described by an equation with one variable controlling for relative thickness (t-value, see [15]). High errors occur in the estimated results from Paleomass in these samples. As for the Atlantic sturgeon (*Acipenser oxyrhynchus oxyrhynchus*), Paleomass fails to bracket its true volume and area with the selected range of k-values (1.6-2.4). But it brackets the hawksbill turtle (*Eretmochelys imbricata*) successfully so that the errors are much lower.

In all the samples of the second test, cross-sectional method has much better performances than GDI or Paleomass, with errors always lower than 2%.

## Discussion

It has been long assumed that the cross-sections of an animal’s main body or limbs can be approximated by ellipses [5]. Based on this assumption some mathematical methods were developed to caluculate the body sizes of animals (e.g., GDI, [13]; mathematical slicing [10]). In some species with rounded or oval cross-sections, these methods do have good performances, as proved in the first test.

In GDI, the semi-major and semi-minor axes of the bases in each slab are measured, then the average values are taken and the slab is treated as a cylinder with elliptical crosssections (see [9] for detailed formula). This assumption is not mathematically rigorous, but it proves to have high accuracy (>95%) when handling objects with near elliptical cross-sections [13]. But errors increase to more than 10% when evaluating irregular-shaped objects (Table 2), revealing the limitation of elliptical approximation.

Motani [11] later argued that biological cross-sections can not always be well-approximated with ellipses. In some cases it is better to use superellipse instead, which is a generalization of ellipse:

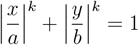

In the first version of Paleomass, Motani [11] used the formula of a superelliptical cone frustum to calculate the volume of each slice (height = 1):

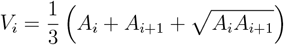

where *A*_*i*_, *A*_*i*+1_ are areas of the two bases. This formula is also not rigorous in math because its validity requires the two bases are similar figures, which is not satisfied in this case. But it still serves well as an approximation formula. This problem is also successfully avoided by the new version of Paleomass implemented in R [15], which invokes vcgVolume() function [25].

The errors of Paleomass in some cases of the two tests may be exaggerated. Its accuracy can be improved by selecting more suitable k-values, but this reveals another weakness of Paleomass. It is not known which k-value range to use when handling animals that have not been examined before. This problem is especially tricky for most extinct animals, as their true body cross-sections are not available. The fin volume and area calculation in Paleomass also presents high errors in the second test. It is possible that a single formula with only one variable controlling for relative thickness may not be sufficient to describe different foils.

Motani [11] noticed that some cross-sections in nature can not be perfectly represented by superellipses. This is supported by the Atlantic sturgeon (*Acipenser oxyrhynchus oxyrhynchus*) model in the second test. It has almost triangular cross-sections (Fig. 3A), and Paleomass fails to bracket its volume and area using k-values from 1.6-2.4 (this range successfully bracketed all models tested in [15]). There are many irregular-shaped cross-sections in nature. It is not possible to list all of them in this paper, but some are presented here as examples (Fig. 3): (1) The cross-sections of fins and limbs in lift-based underwater fliers (e.g., plesiosaurs) and axial swimming tetrapods (e.g., ichthyosaurs and cetaceans) are hydradynamic foils rather than superellipses [26]. (2) Giant salamanders have folds of skin along their body flanks, which lead to irregular-shaped cross-sections. (3) Sea turtles (e.g., leatherback turtle) generally have shells irregular in cross-sections, which can not be approximated using superellipses.

**Figure 3:**
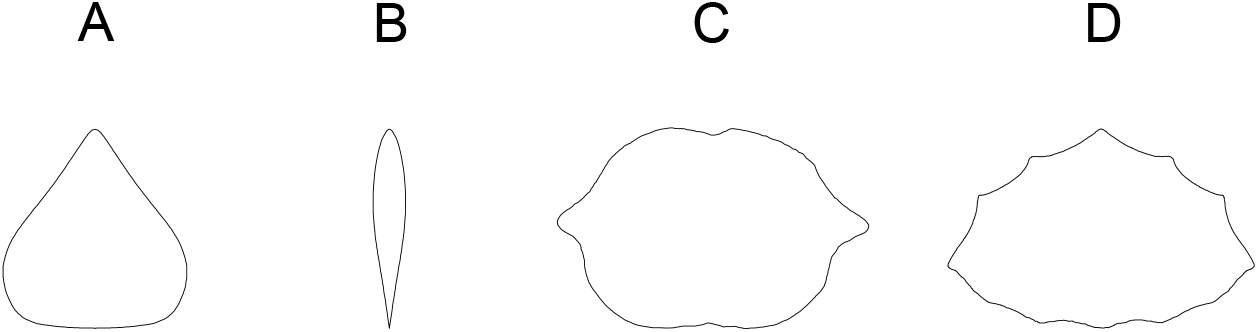
Irregular biological cross-sections. (A) Body cross-section of an Atlantic sturgeon (*Acipenser oxyrhynchus oxyrhynchus*); (B) Cross-section of flipper/fin of secondarily aquatic tetrapods, reproduced from [26]; (C) Body cross-section of a Japanese giant salamander (*Andrias japonicus*); (D) Body cross-sections of a leatherback turtle (*Dermochelys coriacea*). (A)(C)(D) are truncated from accurate 3D models produced by Digital Life within University of Massachusetts at Amherst (downloaded from https://sketchfab.com/DigitalLife3D and used with permission).

The cross-sectional method presented in this study calculates the volume and surface area from cross-sectional profiles directly rather than approximating. This results in more accurate estimation for both volume and area (Table 1, 2). Processing profile images may incorporate extra error, but in all the tests the total errors are lower than 2%. Unlike many previous studies which assume a constant superelliptical k-value (k=2 for ellipse) along the sagittal axis, this method is more flexible by assuming and handling gradually changing cross-sections. It generates point estimation results rather than interval estimations presented by Paleomass, hence the results can be directly incorporated in following studies like scaling regressions (see ‘hybrid approaches’ in [5]). The calculation of cross-sectional method doesn’t require a Cartesian coordinate system, so it is easier to accomplish than some other approaches like polynomial method.

The disadvantage of cross-sectional method is that it requires a series of cross-sectional profiles, which is often unavailable because it didn’t receive enough attention. Cross-sectional outlines can be extracted from front view photos, dissections, 3D scans or precise reconstructions. Paul [27] suggested that accurate skeleton profiles are essential to reconstruct extinct or extant vertebrates, but a rigorous reconstruction of the rib cage is often ignored or not published in previous studies. Careful examination of cross-sections is, however, also suggested in previous studies (e.g., Motani argued that the true cross-sections should be examined when determining k-values [11]). I suggest future scholars pay more attention to detailed and careful reconstruction or acquisition of cross-sectional profiles because simply assuming an elliptical or superelliptical cross-section may lead to high errors, as proved in this paper.

## Conclusion

Cross-sectional method processes cross-sectional profiles directly rather than approximating. It integrates biological cross-sections to calculate volume and surface area. It generates results with a high accuracy, with errors always lower than 2% in all tests. Instead of assuming elliptical or superelliptical cross-sections empirically, future scholars are suggested to carefully examine the profiles to acquire the true shapes.

## Acknowledgement

I thank Beneden Parotodus for assessing math formulas before publication.

